# miR-206 inhibits estrogen-induced proliferation and invasion of ER-α36 positive gastric cancer cells

**DOI:** 10.1101/2021.07.23.453532

**Authors:** Chunyan Yuan, Yuanyuan Yang, Xuebing Jiang, Xiaoli Xie, Jun Liu, Xuming Wang, Xia Sheng

## Abstract

**Objective:** To explore the biological role of miR-206 in inhibiting the proliferation and invasion of estrogen-induced ER-α36 positive gastric cancer cells and its related mechanisms.

**Methods:** MTT and tranwell assays were used to detect the proliferation and invasion of BGC-823 cells under different concentrations and time of ER stimulation. RT-PCR was used to detect the effect of ER stimulation on the content of miR-206 in gastric cancer cells. miR-206 mimics were transfected into BGC-823 cells to detect cell proliferation and invasion; luciferase reporter assay was used to determine whether miR-206 targets the 3’-untranslated region of the CDK14-encoding gene (3’ -UTR).

**Results:** We confirmed that estrogen promoted the proliferation and invasion of BGC-823 cells; As the concentration of estrogen increases, the level of miR206 in the cell gradually rises.Overexpression of miR-206 significantly decreased the estrogen-induced BGC-823 cells. Proliferative and invasive ability; miR-206 inhibits the expression of CDK14 by directly binding to the 3’-UTR of CDK14 mRNA, thereby inhibiting the proliferation and invasion of gastric cancer cells.

**Conclusions:** Our results suggest that miR-206 inhibits estrogen-induced proliferation and invasion of ER-α36-positive gastric cancer cells by inhibiting CDK14 and may be a promising therapeutic option for anticancer therapy.

---

Gastric cancer (GC) is a common type of digestive system cancer. Despite the steady decline in incidence and mortality, the majority of patients are initially diagnosed with late-stage GC. In other words, these patients have already passed up the opportunity to receive radical gastrectomy, the only known cure for GC, at the time of diagnosis [2–4]. This demonstrates the critical need to further our understanding of GC’s pathogenesis and potential molecular mechanisms, which will hopefully aid in the identification of more effective biomarkers and targets for GC diagnosis and treatment [5].

Furthermore, cyclin-dependent kinases (CDKs) are essentially serine-threonine kinases that play an important role in the cell cycle [6]. Due to their regulatory effects on the cell cycle, CDKs have long been thought to be promising targets of anticancer therapy [7]. According to reports, CDK overexpression is present in human cancers such as GC, ovarian cancer, breast cancer, lung cancer and colorectal cancer and is associated with the prognosis of cancer patients [8]. In breast, oesophageal, and GC cells, CDK14, also known as PFTK1 (PFTAIRE protein kinase 1), is moderately to highly expressed [9–11]. Because miRNA-mediated gene regulation is involved in a variety of stem cell mechanisms, miRNA dysregulation is linked to cancer development and metastasis[12,13]. Therefore, it is thought that the presence of regulatory miRNAs is required for the pathogenesis of GC during the differentiation process, where cells change to specialised types. The aforementioned process affects and downregulates the expression of miR-206, which has been shown to have significant effects on cellular differentiation and functions in human rhabdomyosarcoma (RMS) cells. As a result, miR-206 re-expression promotes myogenic differentiation and inhibits tumour growth [14]. Therefore, miR-206 has been identified as a potential target of anticancer therapy [15]. According to breast cancer studies, miR-206 is downregulated in ERα-positive breast cancer cells and directly inhibits ERα expression in MCF-7 cells [16]. Moreover, lower miR-206 expression levels have been linked to late TNM stage, lymph node metastasis, a relatively low total survival rate and increased metastatic potential [17]. However, the underlying molecular mechanisms are not yet fully understood. In this study, it was hypothesised that miR-206 inhibited the proliferation and invasion of GC cells. On this basis, the potential mechanism was discussed, with the goal of providing a reference for clinical treatment of GC.

## Materials and Methods

### 1. Cell culture and transfection

Cell culture and transfection were carried out using a self-owned ER-α36-positive human BGC-823 cell line. BGC-823 cells were cultured in RPMI-1640 containing 10% foetal bovine serum (FBS) plus 100 μg/mL streptomycin at 37°C in a humid incubator with 5% CO_2_. To counteract the effect of oestrogen contained in the medium, RPMI-1640 was replaced with phenol red-free RPMI 1640 supplemented with 5% dextran/charcoal-stripped FBS 24 hours before transfection. Transfection of the cells with miR-206 mimic was accomplished by mixing the transfection reagent Lipofectamine 2000 (Invitrogen, USA) with 100 pmol siRNA oligomer in a reduced serum medium.

### 2. E2 stimulation

For 3 hours, BGC-823 cells were treated with 0.1, 1 and 10 mM E2. Following that, cell viability, migration and invasion rates were all evaluated.

### 3. Cell viability assay

An MTT assay was performed to analyse cell viability in each group. In a 96-well culture plate, a total of 5,000 cells were inoculated. After incubating at 37°C, 5% CO_2_ for 12, 24, 48 and 72 hours, 20 μL of the MTT solution was added to each well (5 mg/ml, Sigma-Aldrich; Merck KGaA, Darmstadt, Germany). Then, the cells were incubated at 37°C for another 4 hours before adding 150 μl dimethyl sulfoxide. After reaction at room temperature for 10 min, the optical density at 570 nm was measured using the Multiskan FC microplate photometer to assess cell proliferation rate.

### 4. Cell invasion assay

Cell invasion was determined using a transwell chamber precoated with Matrigel (BD Biosciences, Franklin Lakes, NJ, USA). A cell suspension (5 × 10^5^ cells / mL) was prepared in DMEM media and subsequently, 300 μL of this cell suspension was added into the upper chamber, and 500 μL of DMEM containing 10 % FBS was added into the lower chamber. After incubation at 37°C in 5% CO_2_ for 24 h, all noninvasive cells were wiped away with a cotton swab. Afterward, 20 μL of MTT solution was added then incubated at 37°C for 4 h. After incubation, 150 μL of DMSO was added and left to incubate for 10 min at room temperature. The cells were then observed and imaged using an inverted microscope.

### 5. Detection by the reverse transcription-quantitative polymerase chain reaction

The TRIzol reagent (Invitrogen; Thermo Fisher Scientific, Inc.) was used to isolate the total RNA from the cells in each group based on manufacturer instructions. To determine miR-206 expression, the miScript Reverse Transcription Kit (Qiagen, Inc., Valencia, CA, USA) was utilised for the reverse transcription of 1 μg total RNA according to manufacturer instructions. Moreover, miScript SYBR Green PCR Kit (Qiagen, Inc.) was used to perform qPCR with ABI 7500 PCR Amplifier (Applied Biosystems; Thermo Fisher Scientific, Inc.). Primers were provided by Guangzhou Fulengen Co., Ltd. (catalogue number HmiRQP9001; Guangzhou, China). The reaction conditions were as follows: 95°C for 5 min and 40 cycles of 95°C for 10 s and 60°C for 30 s. The relative expression of miRNA relative to the relative expression of U6 was standardised by the 2-ΔΔCq method.

### 6. Western blot analysis

The cells were lysed, and a radioimmunoprecipitation assay (RIPA) buffer (Thermo Fisher Scientific, Inc.) was used to separate the protein. The protein concentration was obtained with Bicinchoninic Acid (BCA) Protein Assay Kit (Santa Cruz Biotechnology, Inc., Dallas, TX USA). For Western blot analysis, 60 μg protein was separated with 12% SDS-PAGE gel and transferred to a polyvinylidene fluoride (PVDF) membrane (Thermo Fisher Scientific, Inc.). The PVDF membrane was sealed with 5% skim milk powder in the PBS containing tween (Sigma-Aldrich; Merck KGaA) for 3 h. Afterwards, the membrane, together with mouse anti-CDK14 monoclonal antibody (1:200; AbCAM, Cambridge, MA, USA) or mouse anti-β-actin monoclonal antibody (1:400; AbCAM), was incubated at room temperature for 3 h. The membrane was washed with PBS three times and then incubated with the secondary antibody conjugated with goat anti-mouse horseradish peroxidase (HRP) at room temperature for 1 h. The enhanced chemiluminescence kit (Pierce; Thermo Fisher Scientific, Inc.) was used for chemiluminescence detection (CLD). The protein expression was analysed by the software Image-Pro Plus 6.0 according to the manufacturer’s guide. The β-actin was used as the internal control.

### 7. Luciferase reporter gene assay

Luciferase reporter gene assay was performed to clarify the targeting relation between miR-206 and CDK14 in ER-α-positive gastric cancer (GC) cells. The wild-type (WT) or mutant type (MUT) 3’ untranslated region (UTR) of CDK14 mRNA was inserted downstream the luciferase reporter gene in the pMIR-REPORT vectors. Furthermore, BGC-823 cells were co-transfected with the miR-206 mimics expressing luciferase and the pMIR-REPORT vectors containing WT or MUT CDK14. Afterwards, the cells were incubated in 5% CO_2_ at 37°C for 48 h. The luciferase activity was measured by the Dual-Luciferase Reporter Assay System (Promega Corporation) according to manufacturer instructions.

### 8. Statistical analysis

The data were expressed as mean ± SD of three independent experiments. The statistical software SPSS 17.0 (SPSS, Inc., Chicago, IL, USA) was used for one-way analysis of variance (ANOVA) to analyse the differences between groups. P < 0.05 was considered statistically significant.

## Results

### 1. Stimulation of proliferation and invasion of ER-α36-positive GC cells by oestrogen-signalling

Firstly, the expression levels of ER-α in ER-α36-positive GC cells were detected by cell immunofluorescence assay, the results of which showed that ER-α was highly expressed in ER-α36-positive GC cells (Fig. 1A). Afterwards, the dose-dependent and time-dependent patterns of E2 stimulation on cell proliferation levels in BGC-823 cells were detected by MTT. Additionally, the results showed that with the increase of concentration and time, the proliferation levels of BGC-823 cells increased steadily and reached the maximum under the conditions of 10 mM and 72h (P < 0.001, Fig. 1B). Then, the effect of the treatment with different concentrations of E2 on the invasion of GC cells was detected by the Transwell experiment. Results showed that as concentration increased, the invasion of cells increased and peaked at the concentration of 10 mM (P < 0.001, Fig. 1C). These results indicated that the proliferation and invasion of ER-α36-positive GC cells were stimulated by oestrogen-signalling.

**Figure 1.**
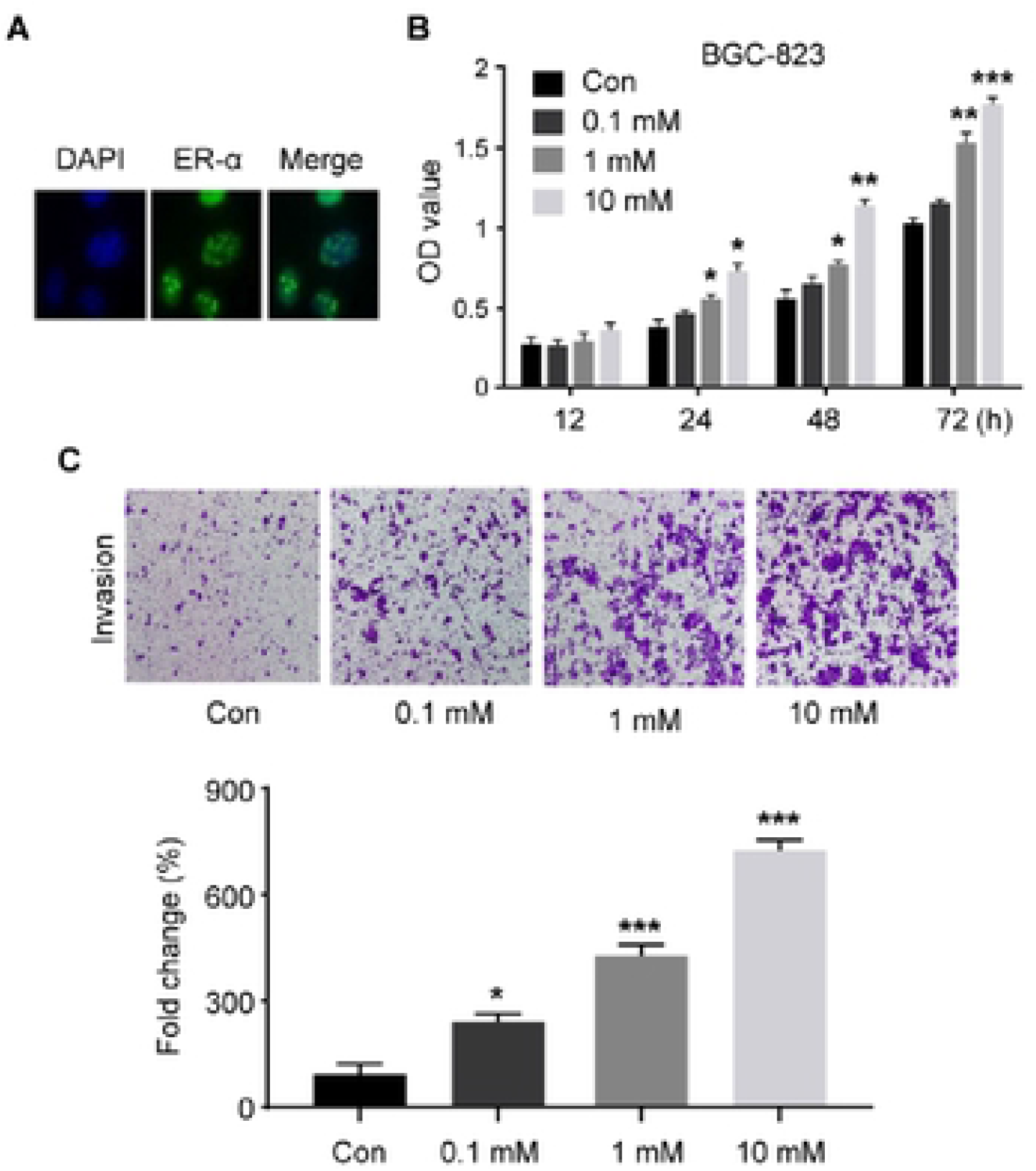
Estrogen signaling stimulates proliferation and invasion of ER-α36-positive gastric cancer cells. A. ER-α is highly expressed in ER-α36 positive gastric cancer cells. B. Dose- and time-dependent patterns of E2 stimulation on cell proliferation levels in BGC-823 cells. C. Effect of different doses of E2 stimulation on the invasive ability of BGC-823 cells. Data are expressed as mean ± SEM of three independent experiments. n=8. *p<0.05, **p<0.01, ***p<0.001 compared with the control group.

### 2. Increase of miR-206 levels in ER-α36-positive GC cells by oestrogen

The effect of the treatment with different E2 concentrations on the miR-206 levels in GC cells was detected by RT-PCR. Results showed that with the increase of concentration, the content of miR-206 in cells increased and peaked at the concentration of 10 mM (P < 0.001, Fig. 2). This indicated that oestrogen increased the miR-206 levels in ER-α36-positive GC cells.

**Figure 2.**
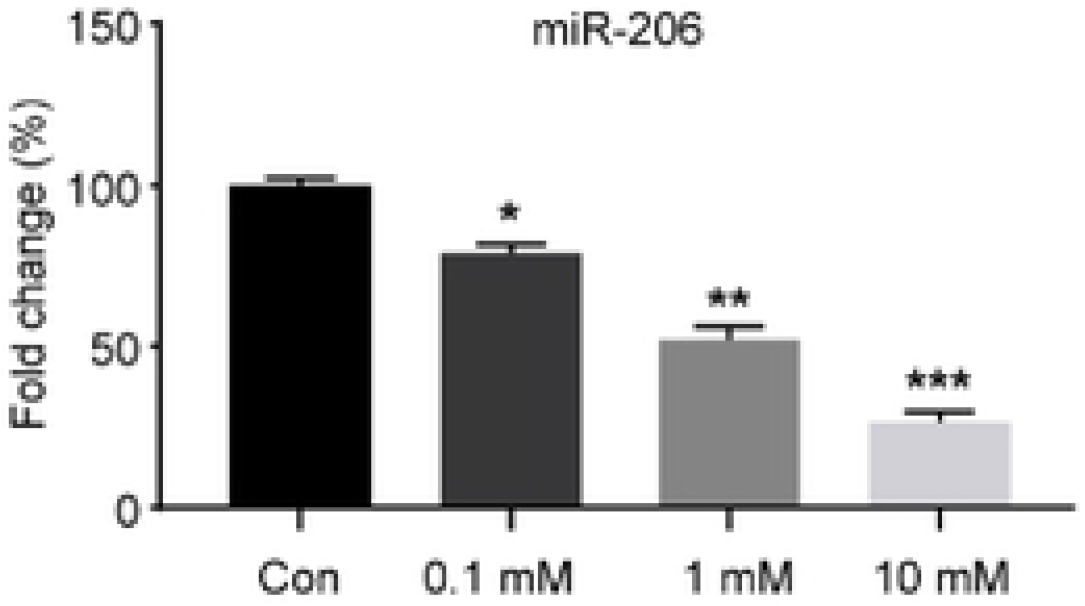
Effect of different concentrations of E2 stimulation on the level of miR-206 in ER-α36 positive gastric cancer cells. Data are expressed as mean ± SEM of three independent experiments. n=8. *p<0.05, **p<0.01, ***p<0.001 compared with the control group.

### 3. Inhibition of oestrogen-induced proliferation and invasion of ER-α36-positive GC cells by miR-206

The present study revealed the effect of miR-206 on the oestrogen-induced proliferation and invasion of ER-α36-positive GC cells. First, the ER-α36-positive GC cells were transfected with miR-206 mimics. The RT-PCR experiment showed that the transfection efficiency could meet the requirements of subsequent experiments (Fig. 3A). Afterwards, compared with the NC group, it was found that the proliferation and invasion of GC cells were significantly induced by oestrogen stimulation, whereas the proliferation and invasion levels were effectively reduced after the transfection with miR-206 mimics. These results indicated that miR-206 could inhibit the oestrogen-induced proliferation and invasion of ER-α36-positive GC cells.

**Figure 3.**
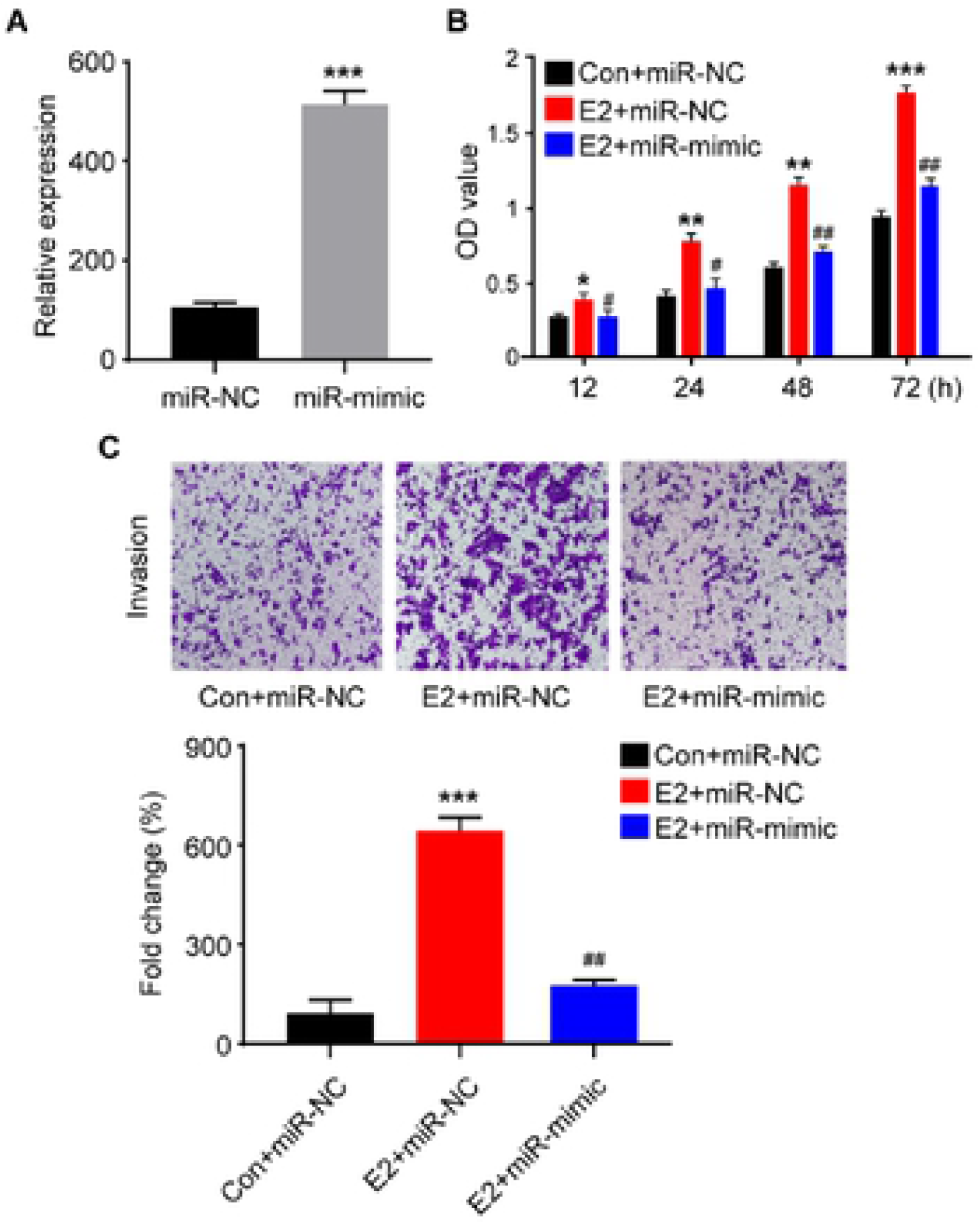
miR-206 inhibits estrogen-induced proliferation and invasion of ER-α36-positive gastric cancer cells. A. RT-PCR to detect the transfection efficiency of miR-206 mimics. B. After transfection of miR-206 mimics in BGC-823 cells, the effect of E2 stimulation on cell proliferation levels was examined. C. After transfection of miR-206 mimics in BGC-823 cells, the effect of E2 stimulation on cell invasion ability was examined. Data are expressed as mean ± SEM of three independent experiments. n=8. *p<0.05, **p<0.01, ***p<0.001 compared with the Con+miR-NC group.

### 4. Exertion of biological effects of miR-206 through specific inhibition of CDK14

Since the proliferation and invasion of GC cells were inhibited by miR-206, the potential mechanism of this effect was studied herein. For this purpose, the online algorithm TargetScan 6.2 was used for bioinformatics analysis in order to predict the potential miRNA targets of miR-206. Among the miRNA targets found to be involved in cell proliferation and invasion, CDK14 was the most favourable. The results of Western blot showed that, under oestrogen stimulation, the expression of CDK14 strongly increased; and with the increase of oestrogen dose, the expression of CDK14 gradually increased (Fig. 4A). TargetScan predicted that the binding sequence in the 3’-UTR of CDK14 is a significant match for miR-206 (Fig. 4B). Moreover, luciferase assay showed that the overexpression of miR-206 inhibited the luciferase activity in cells transfected with WT 3’-UTR plasmid carrying CDK14 gene. In addition, the luciferase activity in MUT CDK14 3’-UTR vectors was not affected by miR-206 transfection (Figs. 4B and C). Therefore, the above data indicated that CDK14 transcripts were the real targets of miR-206.

**Figure 4.**
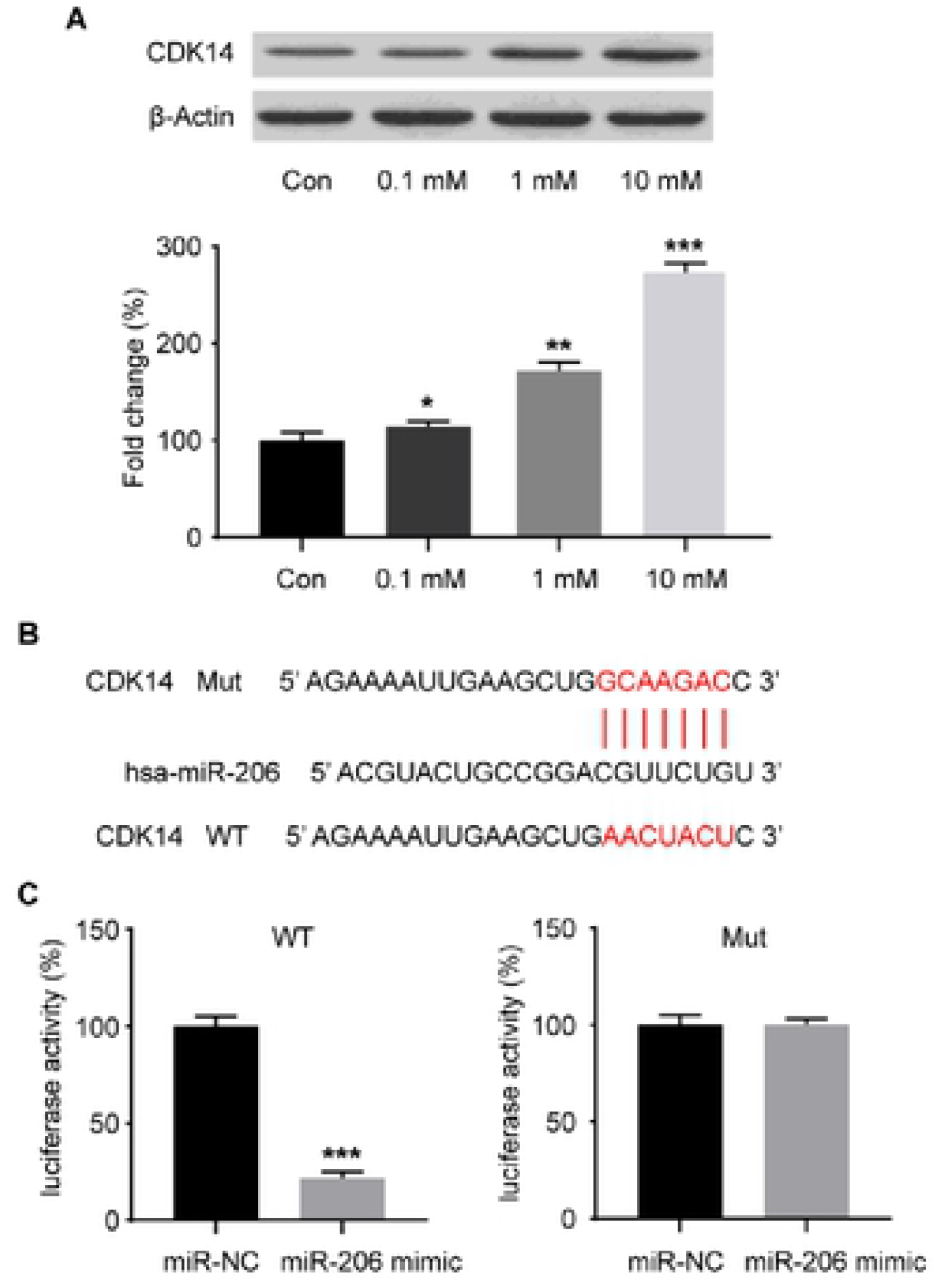
miR-206 inhibits estrogen-induced proliferation and invasion of ER-α36-positive gastric cancer cells by inhibiting CDK14. A. Western blotting assay to detect the expression level of CDK14 after ER-α36 positive gastric cancer cells were stimulated by different concentrations of estrogen. B. Sequence alignment between miR-206 and CDK14. The effect of miR-206 on the 3’-UTR of CDK14 in gastric cancer cells was determined by luciferase activity assay. The cells were co-transfected with the constructed CDK14 wild type (WT) or mutant (MUT) plasmid, miR-206-mimic by lipofectamine 2000. After 24 hours of transfection, the activity of luciferase was measured. The scrambled sequence was used as a negative control (NC). C. Relative value analysis of luciferase reporter experiments. Data are expressed as mean ± SEM of three independent experiments. n=8. *p<0.05, **p<0.01, ***p<0.001 compared with the control group.

## Discussion

The effect of miR-206 on malignant phenotypes of ER-α-positive GC cells was studied herein. It was found that the proliferation and invasion of ER-α-positive GC cells were significantly increased by E2 stimulation, which was accompanied by a decrease in miR-206 levels. The overexpression of miR-206 inhibited the E2-induced upregulation of the proliferation and invasion of GC cells. Further study showed that miR-206 negatively mediated the protein expression of CDK14 by directly binding to the mRNA in GC cells, thus inhibiting the proliferation and invasion of ER-α36-positive GC cells induced by oestrogen.

It has been confirmed that continuous exposure to E2 increases the risk of breast cancer. Moreover, the use of inhibition of E2-mediated signalling for the treatment of oestrogen-dependent GC has been suggested [18,19]. This study revealed that E2 stimulation could significantly increase the viability and invasion of ER-α-positive GC cells. These findings show that few miRs are involved in the development of GC. Further, few miRs have been shown to act as oncogenes or tumour suppressors in GC [20–23]. For instance, it was previously found that miR-939 was significantly downregulated in GC, which was relevant to poor disease-free survival and was considered to be an independent prognostic factor for overall survival [24]. Therefore, expanding the understanding of the miR in GC may contribute to the development of effective strategies for GC treatment.

The progression of tumour metastasis is a continuous process that involves the acquisition of several features through which malignant cells can spread from the primary tumour and colonise at secondary sites. CDK14, a serine/threonine-protein kinase associated with cell division cycle 2, interacts with cyclin D3 and acts as an essential regulator of CDK-cyclins (CCNs) and cell cycle progression [8]. miR-206 may target cyclins; this combination has dramatic effects that further affect cell proliferation and invasion. This suggests that miR-206 is a powerful regulator of proliferation and invasion. The results of this study show that the mechanism of the effect of miR-206 on cell proliferation is relevant to CDK14. Although miR-206 has been reported to activate the apoptosis in HeLa cells [25], apoptosis induced by miR-206 in GC cells was not noted in this study. These data indicate the different tumour suppressive effects of miR-206, including induction of G1 cell cycle arrest and inhibition of cell migration. Therefore, this study showed that miR-206 inhibits the proliferation and invasion in GC cells. Additionally, it was proved that CDK14 is subject to the post-transcriptional regulation by miR-206 and that proliferation and invasion are inhibited by this mechanism. The effects of ER-miR-206-CDK14 were introduced in detail in this study, providing the mechanism for the proliferation and invasion of GC cells and may be used in new therapeutic or diagnostic methods.

## Compliance with ethics guidelines

The experimental protocol was established according to the ethical guidelines of the Helsinki Declaration and was approved by the Human Ethics Committee of Minhang Hospital, Fudan University.

## Authors’ contributions

Chunyan Yuan and Yuanyuan Yang contributed equally to this work.

Chunyan Yuan wrote the manuscript.

Yuanyuan Yang performed literature review and followed-up.

Xia Sheng revised the manuscript.

Xuming Wang revised the manuscript.

All the authors read and approved the final manuscript.

